# The McdAB Carboxysome Positioning System is Widespread Among β-cyanobacteria

**DOI:** 10.1101/737502

**Authors:** Joshua S. MacCready, Joseph L. Basalla, Anthony G. Vecchiarelli

## Abstract

Carboxysomes are protein-based organelles that are essential for allowing cyanobacteria to fix CO_2_. Previously we identified a two-component system, McdAB, responsible for equidistantly positioning carboxysomes in the model cyanobacterium *Synechococcus elongatus* PCC 7942. McdA, a ParA-type ATPase, non-specifically binds the nucleoid in the presence of ATP. McdB, a novel factor that directly binds carboxysomes, displaces McdA from the nucleoid. Removal of McdA from the nucleoid in the vicinity of carboxysomes by McdB causes a global break in McdA symmetry, and carboxysome motion occurs via a Brownian-ratchet based mechanism towards the highest concentration of McdA. Despite the importance for cyanobacteria to properly position their carboxysomes, whether the McdAB system is widespread among cyanobacteria remains an open question. Here, we used neighborhood analysis to show that the McdAB system is widespread among β-cyanobacteria and often clusters near carboxysome-related components. Moreover, we show that two distinct McdAB systems exist in β-cyanobacteria, with Type 2 systems being the most abundant (>98% of β-cyanobacteria) and Type 1 systems, like that of *S. elongatus*, possibly being acquired more recently. Surprisingly, our analysis suggests that the McdAB system is completely absent in α-cyanobacteria. Lastly, all McdB proteins we identified share the sequence signatures of a protein capable of undergoing Liquid-Liquid Phase Separation (LLPS). Indeed, we find that *S. elongatus* McdB undergoes LLPS *in vitro*, the first example of a ParA-type ATPase partner protein exhibiting this behavior. This is an intriguing finding given the recent demonstration of LLPS activity by β-carboxysome core components. Our results have broader implications for understanding carboxysome biogenesis and positioning across all β-cyanobacteria.

**In Brief:** We found that the McdAB carboxysome positioning system is widespread among β-cyanobacteria, absent in α-cyanobacteria, exists in two distinct forms, and that *S. elongatus* McdB undergoes liquid-liquid phase separation.

## Introduction

The ability for cells to organize their interior is ubiquitous across all domains of life. In bacteria, the ParA/MinD family of ATPases have been primarily studied for their ability to segregate genetic cargos, such as chromosomes and plasmids **(Reviews in Baxter and Funnell, 2014; Badrinarayanan et al., 2015).** Less studied are ParA/MinD family members implicated in the positioning of diverse protein complexes, including those involved in secretion **(Viollier et al., 2002; Perez-Cheeks et al., 2012)**, chemotaxis **(Thompson et al., 2006; Ringgaard et al., 2011; Alvarado et al., 2017)**, conjugation **(Atmakuri et al., 2007)**, cell division **(Raskin et al., 1999; MacCready et al., 2017)** and cell motility **(Youderian et al., 2003; Kusumoto et al., 2008)**, as well as protein-based bacterial microcompartments (BMCs), such as the carboxysome **(Savage et al., 2010; MacCready et al., 2018)**. For partitioning plasmids, the ParABS system is the best characterized to date. Mechanistically, ParA proteins dimerize and non-specifically bind DNA in the presence of ATP **(Leonard et al., 2005; Hester and Lutkenhaus, 2007; Castaing et al., 2008; Vecchiarelli et al., 2010)**. Subsequently, ParB, which binds around a centromere-like site on the plasmid, *parS*, displaces ParA from the nucleoid (potentially through ATP hydrolysis stimulation) **(Davis and Austin, 1988; Funnell, 1988; Davis et al., 1992; Bouet and Funnell, 1999; Bouet et al., 2000)**. ParA then recycles its nucleotide and rebinds the nucleoid at a random location. The local formation of ParA depletion zones around individual ParB-bound plasmids results in a global break in ParA symmetry along the nucleoid that ParB-bound plasmids utilize to migrate in a directed and persistent manner towards increased concentrations of ParA on the nucleoid **(Adachi et al., 2006; Hatano et al., 2007; Hwang et al., 2013; Vecchiarelli et al., 2014; Hu et al., 2017)**; a recursive mechanism that ensures equidistant plasmid positioning and faithful plasmid inheritance following cell division.

In our recent study, we identified a new self-organizing ParA-type ATPase system, McdAB, which is responsible for equidistantly positioning the carbon-fixing organelles of cyanobacteria, carboxysomes **(MacCready et al., 2018)**. Carboxysomes are essential for photoautotrophic growth of cyanobacteria. Since O_2_ competes with CO_2_ as a substrate for ribulose-1,5-bisphosphate carboxylase/oxygenase (RuBisCO), the encapsulation of RuBisCO and carbonic anhydrase within a selectively permeable protein shell (the carboxysome) is necessary for generating the high CO_2_ environment needed to drive internal RuBisCO reactions towards the Calvin-Benson-Bassham cycle (CO_2_ substrate) and away from the wasteful process of photorespiration (O_2_ substrate) **(Figure 1AB) (Reviewed in: Kerfeld et al., 2018)**. Comprised of thousands of proteins **(Sun et al., 2019)**, carboxysomes were once thought to be completely paracrystalline **(Kaneko et al., 2006)**. However, recent reports have challenged this idea by showing that carboxysome formation involves the process of Liquid-Liquid Phase Separation (LLPS) **(Wang et al., 2019; Oltrogge et al., 2019)**; a process that describes the ability of proteins to spontaneously demix into dilute and dense phases that resemble liquid droplets **(Reviewed in: Alberti et al., 2019)**. Through these mechanisms, carboxysomes contribute to greater than 25% of global carbon-fixation through atmospheric CO_2_ assimilation **(Rae et al., 2013)**. Using the model rod-shaped cyanobacterium *Synechococcus elongatus* PCC 7942, we identified a small novel protein, McdB, responsible for emergent oscillatory patterning of ATP-bound McdA on the nucleoid **(MacCready et al., 2018)**. While McdB had no identifiable sequence similarities with any known ParB-family members, we found that McdB localized to carboxysomes through multiple shell protein interactions and removed McdA from the nucleoid in their vicinity **(MacCready et al., 2018)**; observations that are analogous to ParA patterning following ParB binding to the *parS* site of plasmids. Thus, like plasmids, we showed that carboxysomes utilize a Brownian-ratchet based mechanism whereby McdB-bound carboxysome motion occurs in a directed and persistent manner towards increased concentrations of McdA on the nucleoid **(Vecchiarelli et al., 2014; Hu et al., 2017; MacCready et al., 2018) (Figure 1C)**.

**Figure 1:**
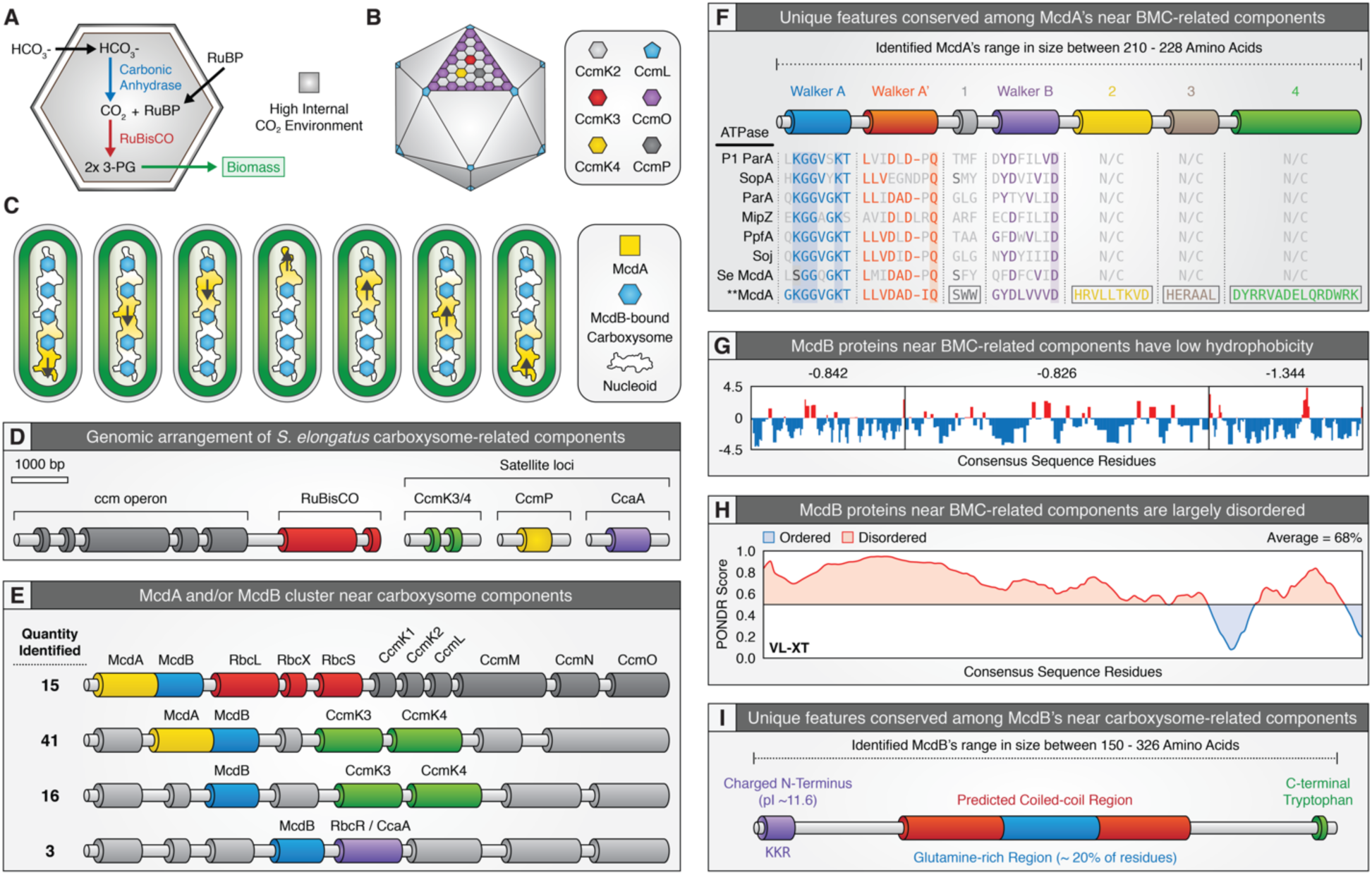
Candidate McdA/B proteins cluster near known carboxysome components and share common unique features. **(A)** Illustration of internal carboxysome enzymatic reactions. **(B)** Illustration of carboxysome protein shell. **(C)** Individual McdB-bound carboxysomes move towards increased concentrations of McdA on the nucleoid that drives equal spacing. **(D)** Representative illustration showing that carboxysome-related genes are found across multiple loci in *S. elongatus*. **(E)** Representative illustration of the genomic context of McdA and/or McdB near carboxysome-related components. **(F)** Conserved features among McdA proteins found near carboxysome components. Known conserved ParA regions, deviant-Walker A (blue), A’ (red) and B (purple) boxes, are conserved among all classic ParA proteins, *S. elongatus* McdA, and putative McdA proteins identified near carboxysome components (* * McdA). Classic ParA ATPase proteins shown: *Escherichia coli* phage P1 ParA (plasmid partitioning - YP_006528), *Escherichia coli* (strain K12) F plasmid SopA (plasmid partitioning - NP_061425), *Caulobacter crescentus* ParA (chromosome segregation - AAB51267), *Caulobacter crescentus* MipZ (chromosome segregation - NP_420968), *Rhodobacter sphaeroides* PpfA (chemotaxis distribution - EGJ21499) and *Bacillus subtilis* Soj (chromosome segregation - NP_391977). Regions conserved among only McdA proteins found near carboxysomes components: Double tryptophan region 1 (grey), and regions 2 (yellow), 3 (brown) and 4 (green). **(G)** Consensus amino acid sequence from identified McdB proteins exhibit a low hydrophobicity. **(H)** McdB proteins are predicted to be intrinsically disordered. **(I)** Conserved features among McdB proteins found near carboxysome components. Charged N-terminal domain (purple), predicted coiled-coil (red), glutamine-rich region within coiled-coil (blue) and C-terminal tryptophan residue (green).

Several important questions remained following our study. First, it was unclear whether the McdAB system was widespread among cyanobacteria, especially given that cyanobacteria are an incredibly diverse and widely distributed phylum of bacteria that display complex morphologies, including: (i) unicellular, (ii) baeocystous, (iii) filamentous, (iv) heterocystous, and (v) ramified. Cyanobacteria are also taxonomically classified as α or β depending on the form of RuBisCO they encapsulate; β-cyanobacteria encapsulate form 1B RuBisCO in β-carboxysomes and α-cyanobacteria encapsulate form 1A RuBisCO in α-carboxysomes **(Rae et al., 2013)**. These two types of carboxysomes are structurally distinct, and α-carboxysomes are thought to have been horizontally transferred to cyanobacteria from chemoautotrophic Proteobacteria **(Rae et al., 2013)**. Thus, it’s not obvious whether α-cyanobacteria possess a McdAB system and whether it would share similarities to that of the β-cyanobacterial McdAB system. Second, we noted in our previous study that McdA surprisingly lacked the signature lysine residue in the Walker A ATP-binding motif – a critical residue that defines the ParA family of ATPases. While BLASTp results for McdA identified additional McdA-like sequences in other cyanobacteria that also lacked this lysine residue, the results were few and many were plasmid encoded. Therefore, it was not clear whether the McdAB system was unique to *S. elongatus* or if more than one type of McdAB system evolved in cyanobacteria. Lastly, it was not obvious why reliable results from BLASTp for *S. elongatus* McdB could not be obtained. Interestingly however, our neighborhood analysis around the carboxysome operon in the distantly-related cyanobacterium *Gloeobacter kilaueensis* JS1 identified a ParA-type ATPase that possessed the signature lysine residue absent in *S. elongatus* McdA and a small downstream coding sequence. The protein product of this gene loaded onto *S. elongatus* carboxysomes, which was surprising because it had no sequence homology to *S. elongatus* McdB, **(Figure 8AB: MacCready et al., 2018)**. The findings suggest a more rigorous gene neighborhood analysis was necessary to identify other McdAB systems.

Here, we performed a neighborhood analysis for McdA/B-like sequences encoded near carboxysome components in 537 cyanobacterial genomes (205 α-cyanobacterial and 332 β-cyanobacterial). Our analysis revealed that the McdA/B system is widespread among β-cyanobacteria and is surprisingly absent in α-cyanobacteria. Across these β-cyanobacteria, McdA/B were found near carboxysome components in ∼31% of genomes and near the minor shell components CcmK3 and CcmK4 in ∼25% of genomes; suggesting a strong functional association. Our analysis also shows that there are two types of McdAB systems, which we term type 1 & 2. Type 1 systems, like that of *S. elongatus*, consist of a McdA without the signature lysine residue and a McdB with a predicted C-terminal coiled-coil. Type 2 systems alternatively, which were found to be the most abundant (>98% of genomes), consist of a McdA ATPase with the signature lysine residue present, and a McdB with a predicted central coiled-coil. The low representation of the *S. elongatus* Type 1 McdAB system suggested a possible unique origin. In support of this hypothesis, our cyanobacterial phylogeny strongly suggests that *S. elongatus* is immediately adjacent to α-cyanobacteria, which lack the McdAB system, and shares a common ancestor with cyanobacteria where a McdAB system could not be identified. Lastly, when comparing identified McdB proteins, little to no sequence homology was observed. However, all shared the known hallmarks of proteins capable of LLPS: (i) intrinsic disorder, (ii) biased amino acid compositions, (iii) low hydrophobicity, and (iv) extreme multivalency. We purified *S. elongatus* McdB and found that it forms phase-separated droplets across a physiological pH-range. To our knowledge, this is the first experimental demonstration of LLPS behavior in a ParA-type ATPase partner protein and is an interesting finding given that carboxysome formation has recently been shown to involve LLPS **(Oltrogge et al., 2019; Wang et al., 2019)**. Collectively, these results have broad implications for understanding carboxysome formation, homeostasis, positioning, and function.

## Results

### Finding Homologs of S. elongatus McdA/B

To explore whether the McdAB system is widespread among cyanobacteria, we began our analysis by performing BLASTp searches for *S. elongatus* McdA (Synpcc7942_1833) and McdB (Synpcc7942_1834). We previously reported that the deviant Walker A box of *S. elongatu*s McdA lacks the signature lysine residue that defines the ParA family of ATPases (**K**GGXXKS/T) **(MacCready et al., 2018)**. The serine substitution at this position in McdA (**S**GGQGKT) may underlie the unusually high ATPase activity of McdA, which displays a maximum specific activity that is roughly two-orders of magnitude greater than that of other well-studied ParA systems **(Ah-Seng et al., 2009; Vecchiarelli et al., 2010; MacCready et al., 2018)**. Our BLASTp search results for *S. elongatus* McdA returned only a few McdA-like sequences where the signature lysine residue was replaced with a serine **(Figure S1A)**. Four of these hits were nearly identical to *S. elongatus* McdA (*Synechococcus elongatus* PCC 6301, *Synechococcus elongatus* PCC 11801, *Synechococcus elongatus* UTEX 3055, and *Synechococcus sp.* UTEX 2973). Recent crystallization of a plasmid-encoded McdA-like protein from the cyanobacterium *Cyanothece sp.* PCC 7424 (PCC7424_5529) showed that a lysine residue in the middle of the protein (K151) functioned analogously to the signature lysine residue of classical ParA-type ATPases found within the Walker A box **(Figure S1A)**. K151 was found to contact the oxygen between the β and γ phosphates of ATP and promote formation of a sandwich dimer **(Schumacher et al., 2019)**. Consistent with this finding, our McdA-like BLASTp hits also possessed the K151 residue **(Figure S1A)**.

BLASTp results for *S. elongatus* McdB were extremely poor. However, when performing a gene neighborhood analysis of the newly identified McdA-like sequences above, a short open reading frame was identified immediately downstream. While these sequences shared high similarity amongst themselves, they largely differed from *S. elongatus* McdB outside the first ∼25 amino acids **(Figure S1B)**. ParB proteins possess a small charged region at their N-terminus responsible for interacting with its cognate ParA protein and stimulating its ATPase activity **(Radnedge et al., 1998; Ravin et al., 2003; Barillà et al., 2007; Ah-Seng et al., 2009)**. Consistent with this observation, the recent crystallization and analysis of the *Cyanothece sp.* PCC 7424 McdB-like protein (PCC7424_5530) revealed that the N-terminus (AA 1-150) mediated interaction with the *Cyanothece sp.* PCC 7424 McdA-like protein **(Schumacher et al., 2019)**. Given the similarity between *S. elongatus* McdA and the McdA-like sequences we identified by BLASTp, it is not surprising that the N-terminal region of *S. elongatus* McdB and the McdB-like sequences identified by BLASTp were highly conserved in this region **(Figure S1B)**. However, outside this N-terminal region, it wasn’t obvious why *S. elongatus* McdB greatly differed from our newly identified McdB-like sequences. The structure of the McdB-like protein from *Cyanothece sp.* PCC 7424 possessed two small central helices followed by a large C-terminal coiled-coil region **(Schumacher et al., 2019)**. While, *S. elongatus* McdB is predicted to also possess a C-terminal coiled-coil **(MacCready et al., 2018)**, *S. elongatus* McdB possesses a large glutamine-rich region and several extensions and gaps relative to the McdB-like BLASTp hits **(Figure S1B)**; potentially due to recognition of different cargos (i.e. carboxysomes vs. plasmids). Indeed, many of these McdAB-like proteins were plasmid encoded and were also the sole ParA-type system on these plasmids. Collectively, given that: (i) several of our McdA/B hits were plasmid encoded, suggesting that these proteins might function in plasmid partitioning instead of carboxysome positioning, (ii) only a few homologs were identified among the hundreds of available cyanobacterial genomes, (iii) the McdB-like protein in *G. kilaueensis* JS1 that we previously showed loaded onto *S. elongatus* carboxysomes **(MacCready et al., 2018)** followed a ParA-type ATPase that possessed the signature lysine residue and lacked K151, and (iv) *G. kilaueensis* JS1 McdAB were encoded within the carboxysome operon, we reasoned that these proteins identified by BLASTp above were not true McdAB carboxysome positioning systems. Therefore, an alternative more rigorous analysis was needed to identify McdAB homologs across cyanobacteria.

### McdA/B Co-occur with Carboxysome Components in Many Cyanobacteria

The increased availability of genomic data, in combination with targeted bioinformatic analyses, has resulted in a wealth of data suggesting that the vast majority of BMC-related genes tend to form operons with their respective encapsulated enzymes. For example, the most recent bioinformatic efforts to identify novel BMCs among bacteria resulted in the identification of 23 different types of BMCs across 23 different bacterial phyla **(Axen et al., 2014)**. This study showed that neighborhood analyses are a powerful and reliable tool for the identification of new factors involved with BMC function. Therefore, since BLASTp was an unreliable method to identify new candidate McdA and McdB proteins, we next performed neighborhood analysis for McdAB-like sequences that clustered near carboxysome-related components across 537 cyanobacterial genomes (205 α-cyanobacterial and 332 β-cyanobacterial). We defined candidate McdA sequences as having a deviant Walker A motif with global homology to ParA-type ATPases and candidate McdB sequences as the protein product of the open reading frame immediately following these McdA sequences. We note that additional carboxysome-related genes are often located at distant loci from the main *ccm* (Carbon Concentrating Mechanism) operon **(Figure 1D) (Axen et al., 2014; Sommer et al., 2017)**.

Using these criteria, our analysis identified 75 examples of McdA/B clustering near carboxysome components (genomic distances of 2209 bp ± 1679 bp or 2 genes ± 1.6 genes upstream or downstream of the carboxysome component). Of these, McdAB-like sequences clustered near the *ccm* operon in 15 species and near the distant loci of the minor shell proteins CcmK3 and CcmK4 in 41 species **(Figure 1E)**. Moreover, in 16 species, only a McdB-like sequence clustered with CcmK3 and CcmK4, and in 3 species, a McdB-like sequence was found near either the RuBisCO transcription factor (RbcR) or carbonic anhydrase (CcaA) **(Figure 1E)**. In both these instances, a McdA-like sequence was not identified, suggesting McdB proteins in general may have additional functions independent of its role in positioning carboxysomes via interaction with McdA.

To identify additional McdA and McdB proteins among cyanobacteria that did not cluster near carboxysome components, we needed to establish a new criterion to aid our search. To accomplish this, we generated multiple sequence alignments for the McdA and McdB proteins that clustered near carboxysome components to identify highly conserved regions and/or features among these proteins **(Figure S2AB)**. Among the 48 McdA proteins, the first notable features we found were that they could range in size from 210-228 amino acids and that each of these proteins possessed the signature lysine residue within the Walker A motif, unlike that of *S. elongatus* McdA which instead possesses a serine residue **(Figure 1F and Figure S2A) (MacCready et al., 2018)**. Secondly, two additional threonine residues followed the last threonine residue of the Walker A motif **(Figure S2A)**. Lastly, two tryptophan residues adjacent to the Walker A’ motif (1 – grey cylinder) and three additional regions towards the last half of these proteins (2 – yellow cylinder, 3 – brown cylinder and 4 – green cylinder) were present among all McdA proteins encoded near carboxysome components, but not among classical ParA proteins or *S. elongatus* McdA **(Figure 1F and Figure S2A)**.

Unlike these McdA proteins, conservation among McdB proteins near carboxysome components was extremely low; partly due to the surprising observation that McdB proteins ranged in size from 150-326 amino acids. However, several shared features existed, which allowed us to establish criteria to define an McdB protein. First, we found that all McdB proteins were largely polar and biased in amino acid composition **(Figure 1G)**. Second, Predictor of Natural Disordered Regions (PONDR) predicted McdB proteins to be highly disordered (Average disorder = 68%) **(Figure 1H) (Romero et al., 2001)**, which explains our poor BLASTp results here and prior **(MacCready et al., 2018)**. Lastly, all identified McdB proteins possessed a highly charged N-terminal region, a predicted coiled-coil, a glutamine-rich region centrally-located within the predicted coiled-coils, and a tryptophan residue within the last 4 residues of each sequence **(Figure 1I and Figure S2B)**. The bioinformatics analysis suggests all McdB proteins we identified share known hallmarks of phase-separating proteins: largely polar, low amino acid complexity, low hydrophobicity, and intrinsic disorder.

### The McdAB System is Widespread Among β-cyanobacteria

Using our new criteria for McdAB proteins established from McdAB proteins that cluster near carboxysome components, we performed an additional manual search within α- and β-cyanobacterial genomes where McdAB proteins were not previously identified. This approach was necessary given that cyanobacteria possess enormously diverse genomic architectures and relatively poor operon structure in comparison to other well-studied bacteria **(Beck et al., 2018)**. Not a single McdAB system that fit within our criteria were identified among the 205 α-cyanobacterial genomes we analyzed. Intriguingly, many α-cyanobacteria did not possess a single homolog of a ParA-type ATPase family member in general. But in β-cyanobacteria, we identified several McdA and McdB sequences that fit within our criteria. In total, we found 248 McdA (∼75% of β-cyanobacterial genomes) and 285 McdB sequences (∼86% β-cyanobacterial genomes) from 332 β-cyanobacterial genomes analyzed **(Figure 2A and Table 1)**. We note that 16% of β-cyanobacterial genomes without candidate McdA, 8% of β-cyanobacterial genomes without McdB, and 9% of β-cyanobacterial genomes without McdA and McdB were not fully assembled genomes (still in scaffolds or contigs). Collectively, we found that McdA proteins ranged in lengths from 202-253 amino acids, while the lengths of McdB proteins varied far greater, ranging in from 132-394 amino acids. In 199 β-cyanobacterial genomes, McdAB were found next to each other genomically (∼82% of β-cyanobacterial genomes with McdAB) **(Figure 2A)**. Moreover, McdA and/or McdB clustered near carboxysome-related components in 75 genomes (∼31% of β-cyanobacterial genomes with McdAB) and clustered near the distant locus for the minor shell components CcmK3 and CcmK4 in 60 genomes (∼25% of β-cyanobacterial genomes with McdAB). This finding suggests a strong functional association between McdB and CcmK3/CcmK4, which is consistent with our previous bacterial two-hybrid results showing a strong interaction between McdB and these two shell components of the carboxysome **(Figure 3F: MacCready et al., 2018)**. Lastly, McdAB sequences were found not only among all major taxonomic orders of β-cyanobacteria **(Figure 2B)** but were also found in genomes across all 5 major morphologies **(Figure 2C)**.

**Figure 2:**
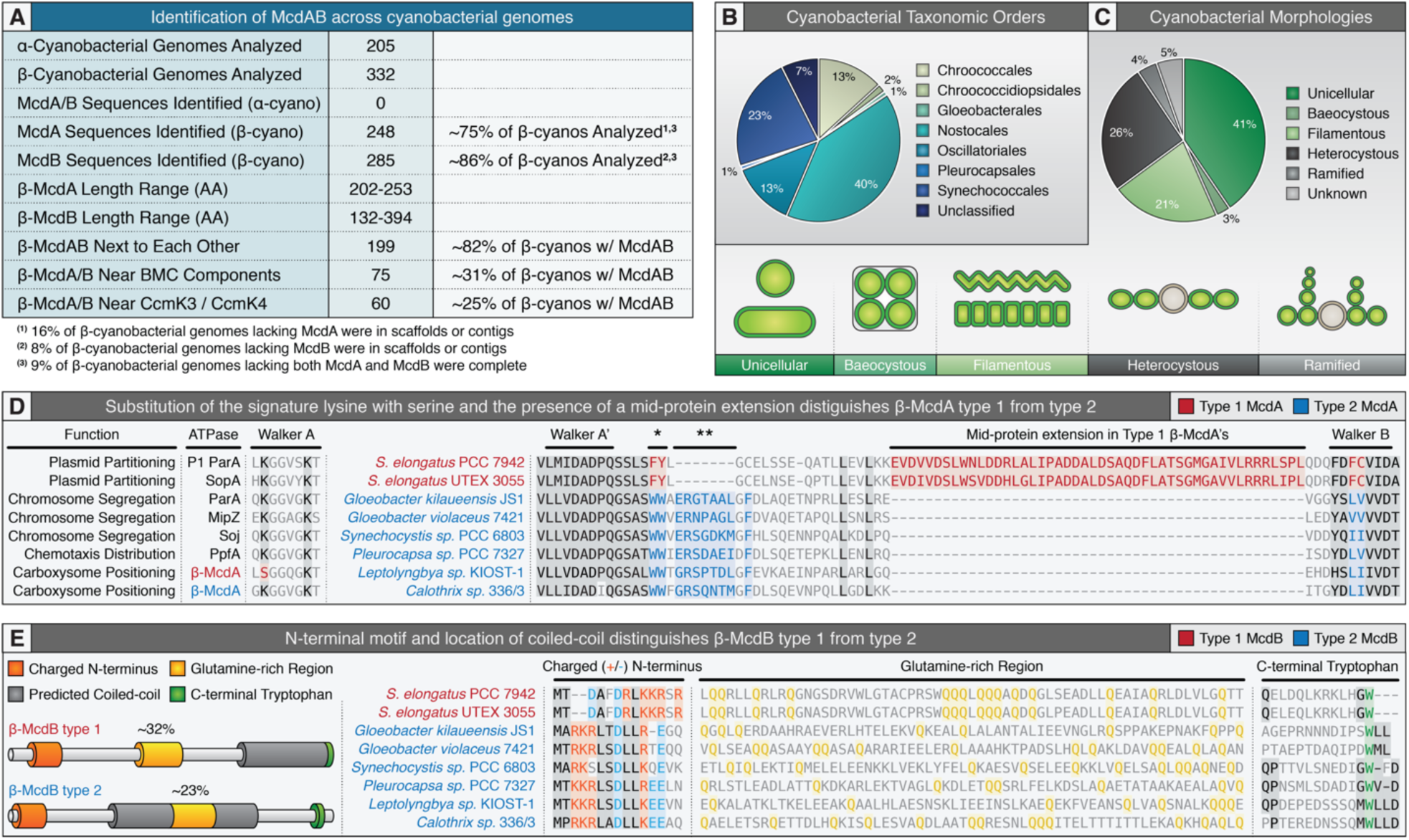
Two distinct McdAB systems exist in β-cyanobacteria. **(A)** Table highlighting the prevalence of certain sequence features for all McdA/B proteins identified among cyanobacteria. **(B)** McdA/B are widely distributed among cyanobacterial taxonomic orders. **(C)** McdA/B are found in all 5 major morphologies of cyanobacteria. General illustration of cyanobacterial morphologies below. **(D)** Type 1 McdA proteins (red) are distinct from Type 2 McdA proteins (blue). Left: Type 1 McdA proteins (red) possess a serine instead of lysine in the Walker A box. Type 2 McdA proteins (blue) possess the signature lysine. Middle: Areas of conservation are shaded black. Conserved regions unique to Type 1 McdA proteins shaded red. Conserved regions unique to Type 2 McdA proteins shaded blue. The Walker A’ box is conserved among both McdA types. Type 2 McdA proteins have a highly conserved double tryptophan region (*) not found in Type 1. Type 2 McdA proteins have a small 7 amino acid insertion (* *) and highly conserved phenylalanine, glutamic acid and proline residues following the double tryptophan region. Type 1 McdA proteins have a large internal extension not found in Type 2. Right: Walker B box is generally conserved, but Type 1 McdA proteins possess phenylalanine and cysteine residues that are instead small polar residues in Type 2 McdA proteins. McdA sequences shown: *Synechococcus elongatus* PCC 7942 (Synpcc7942_1833), *Synechococcus elongatus* UTEX 3055 (Unannotated), *Gloeobacter kilaueensis* JS1 (GKIL_0670), *Gloeobacter violaceus* 7421 (glr2463), *Synechocystis sp.* PCC 6803 (MYO_127120), *Pleurocapsa sp.* PCC 7327 (Ple7327_2492), *Leptolyngbya sp.* KIOST-1 (WP_081972678), and *Calothrix sp. 336/3* (AKG24853). **(E)** Type 1 McdB proteins (red) are distinct from Type 2 McdB proteins (blue). Left: Type 1 McdB proteins have a charged N-terminal domain (orange), central glutamine-rich region (yellow), C-terminal coiled-coil (grey) and terminal tryptophan residue (green). Type 2 McdB proteins have a charged N-terminal domain (orange), central coiled-coil (grey), glutamine-rich regions within the coiled-coil (yellow) and a C-terminal tryptophan within the last four amino acids. Middle: The N-terminal charged (positive charge - orange, negative charge - blue) domain is inverted between Type 1 and Type 2 McdB proteins. Middle: Glutamine-rich regions (yellow). Right: All McdB proteins have a tryptophan residue within the last four amino acids (green). McdB sequences shown: *Synechococcus elongatus* PCC 7942 (Synpcc7942_1834), *Synechococcus elongatus* UTEX 3055 (Unannotated), *Gloeobacter kilaueensis* JS1 (GKIL_0671), *Gloeobacter violaceus* 7421 (glr2464), *Synechocystis sp.* PCC 6803 (MYO_127130), *Pleurocapsa sp.* PCC 7327 (Ple7327_2493), *Leptolyngbya sp.* KIOST-1 (WP_035984653), and *Calothrix sp.* 336/3 (Unannotated).

**Figure 3:**
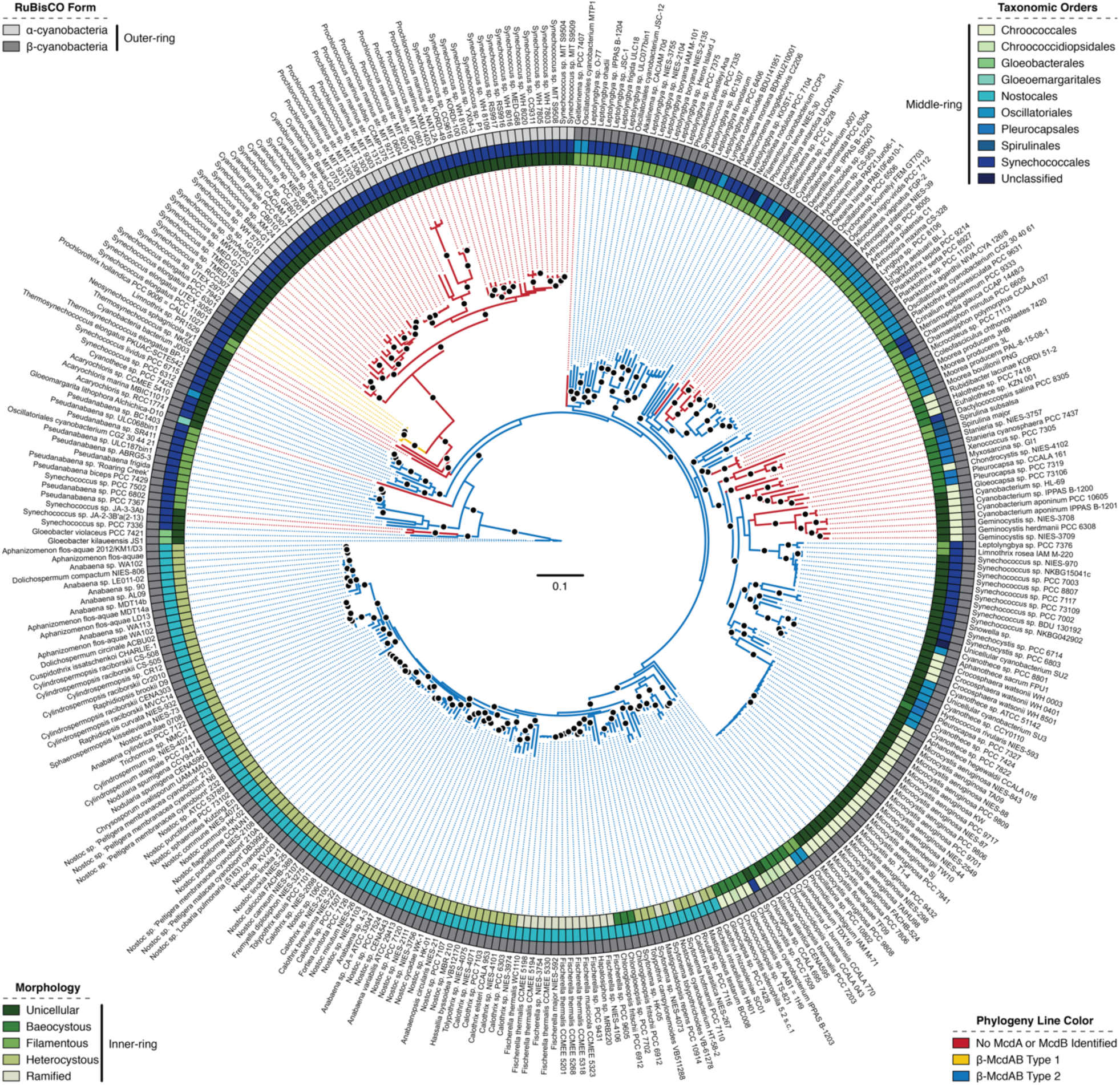
A possible unique origin for the Type 1 McdAB system. Inferred cyanobacterial phylogeny of genomes analyzed. Outer ring: cyanobacterial RuBisCO type. Middle ring: cyanobacterial taxonomic order. Inner ring: cyanobacterial morphology. Line color: Type 1 McdAB systems (yellow), Type 2 McdAB system (blue) and no identified McdAB system (red). Black dot represents > 70% support (500 replicates).

### Two Distinct McdAB Systems Exist in β-cyanobacteria

While conducting this bioinformatic analysis, we found that the vast majority (∼98%) of newly identified McdAB proteins were quite distinct from *S. elongatus* McdAB and only four additional cyanobacterial species, which are closely related to *S. elongatus*, had a similar McdAB system to that of *S. elongatus*. Therefore, we designated the two McdAB systems as Type 1, those similar to *S. elongatus* McdAB, and Type 2, those similar to the newly identified McdAB’s in this study **(Figure 2DE)**. Several features distinguish McdA as Type 1 or Type 2. For example, Type 1 McdA proteins lack the signature lysine residue within the Walker A motif, whereas Type 2 McdA’s possess this lysine residue like classical ParA family members. The double tryptophan found in all Type 2 McdA proteins (*) is instead a phenylalanine and a tyrosine in Type 1 McdAs **(Figure 2D)**. Additionally, a small ∼ 7 amino acid insertion (* *) immediately following the double tryptophan as well as a downstream phenylalanine are highly conserved in Type 2 McdA’s, but absent in Type 1 **(Figure 2D)**. Most notably, Type 1 McdA’s have a long extension between the Walker A’ and Walker B motifs that is not present in Type 2 McdA’s **(Figure 2D)**. Lastly, the Walker B motif of Type 1 McdA’s has a conserved phenylalanine and cysteine that are instead small hydrophobic residues in Type 2 McdA’s.

We also found the two types of McdB significantly differed. For example, Type 1 McdB’s have a predicted coiled-coil at the C-terminus, while all Type 2 McdB’s have a predicted coiled-coil near the middle of the protein **(Figure 2E)**. Moreover, the charged N-terminal region appeared inverted among Type 1 and Type 2 McdB’s, where negatively charged residues proceeded positively charged residues in Type 1 McdB’s and positively charged residues proceeded negatively charged residues in Type 2 **(Figure 2E)**. Interestingly, although both types have a central glutamine-rich region, Type 1 McdB’s are almost 10% more enriched for glutamine than Type 2 McdB’s **(Figure 2E)**. Despite these differences, we found that both types possessed an invariant C-terminal tryptophan **(Figure 2E)**. Taken together, these features strongly suggest that two distinct McdAB systems are present among cyanobacteria and that the Type 2 system identified in this study is the most widely conserved.

### A Unique Origin for Type 1 McdAB Systems?

Since we found that α-cyanobacteria lack McdAB and that Type 1 systems were only present in ∼2% of cyanobacteria studied, we next sought to better understand the phylogenetic placement of Type 1 and Type 2 McdAB systems and determine whether the Type 1 McdAB system could have evolved independent of Type 2 systems. To explore this, we generated a Maximum-Likelihood tree inferred using a concatenation of the proteins DnaG, RplA, RplB, RplC, RplD and RplE, which have recently been shown to be good-markers for cyanobacterial phylogenetic reconstruction **(Hirose et al., 2019)**. Notably, we found that the Type 2 McdAB system was widespread among cyanobacteria including our outgroup taxonomic order Gloeobacterales, considered the “most primitive” order among living cyanobacteria due to a lack of thylakoid membranes **(Rippka et al., 1974; Guglielmi et al., 1981; Nakamura et al., 2003)**. This was an important finding that suggested the Type 2 McdAB system was present at the earliest known point of cyanobacterial evolution. Alternatively, the Type 1 McdAB system was only present in cyanobacterial species adjacent to α-cyanobacteria, which we found to lack the McdAB system, and shared a common ancestor with four β-cyanobacterial species where we could not identify an McdAB system of either type **(Figure 3)**. This was a surprising result that suggested a more recent origin for the Type 1 McdAB system. Our phylogenic inference also suggested multiple independent losses of the McdAB system with no obvious shared characteristics among species (i.e. different taxonomic orders and morphologies) **(Figure 3)**. Together, these results suggest that the Type 2 McdAB system is more ancestral and that Type 1 evolved independently of the Type 2 McdAB system, possibly via horizontal gene transfer.

### McdB Proteins Possess the Hallmarks for Phase Separation

Why McdB proteins were so highly diverged in primary sequence was unclear. For example, while Type 2 McdA’s have high mean amino acid similarity (∼64%), we found that the mean amino acid similarity among Type 2 McdB’s was extremely low (∼20%) **(Figure 4AB)**. Partially contributing to this diversity among McdB proteins, we found that the lengths of the central coiled-coils and the N- and C-terminal extensions from the coiled-coil greatly varied in length **(Figure 4CDE)**. N-terminal extensions were 60 ± 23 amino acids, central coiled-coils were 93 ± 25 amino acids, and the C-terminal extensions were 56 ± 16 amino acids in length **(Figure 4CDE)**. ProtScale prediction **(Scale: Kyte and Doolittle, 1982)** determined that both Type 1 and Type 2 McdB’s were largely hydrophilic, and that Type 1 McdB’s possessed a small hydrophobic patch towards the middle of the protein **(Figure 4F)**. Consistent with this finding, all three Type 2 McdB domains were biased towards hydrophilic amino acids and had obvious repetitive sequences that were rarely similar from one McdB protein to the next **(Figure 4G)**.

**Figure 4:**
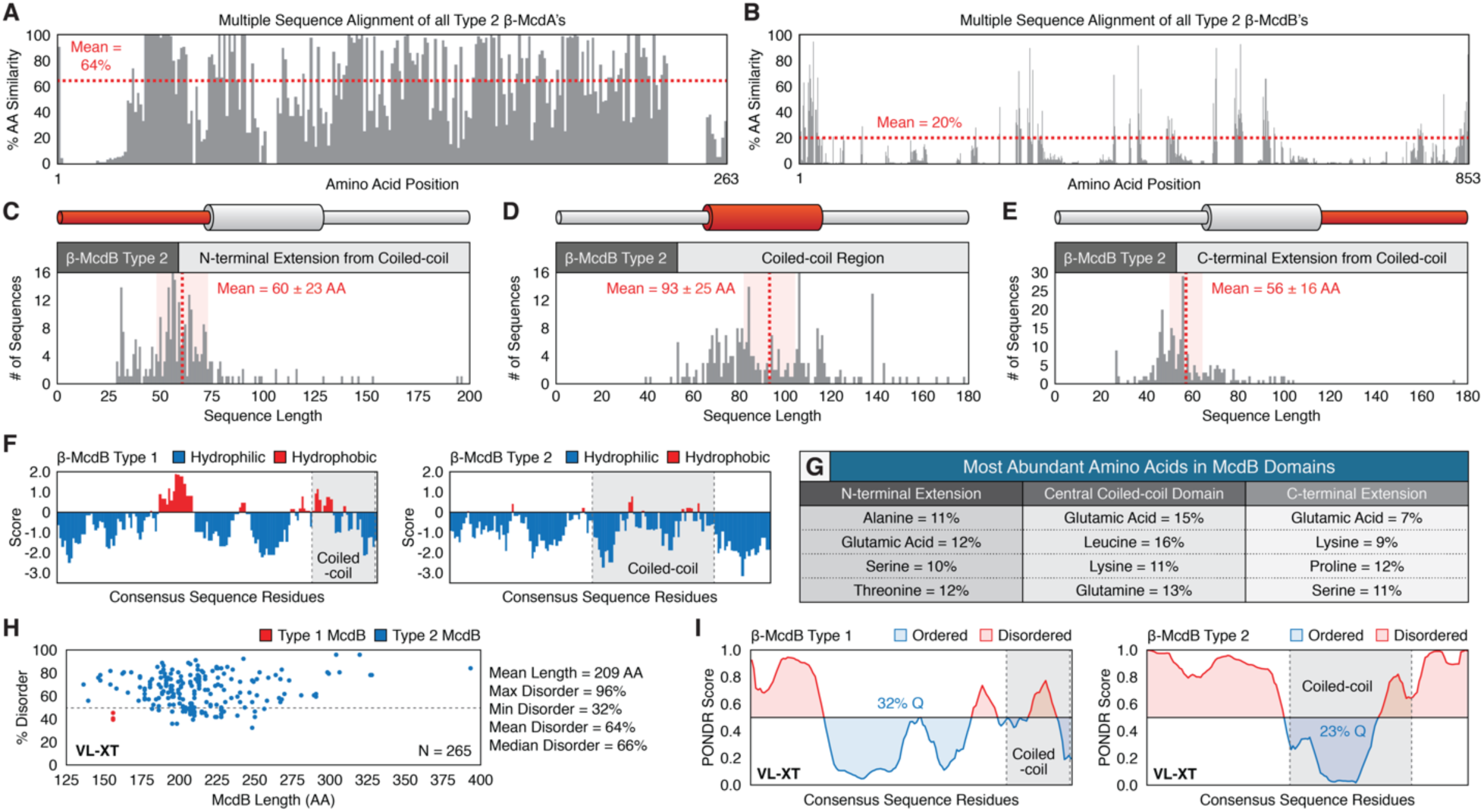
Two distinct McdAB systems exist in β-cyanobacteria. **(A)** Plot of amino acid similarity from multiple sequence alignment of Type 2 McdA proteins (n=XX sequences). **(B)** Plot of amino acid similarity from multiple sequence alignment of Type 2 McdB proteins (n=XX sequences). **(C)** Plot of Type 2 McdB N-terminal extension lengths, **(D)** coiled-coil lengths, and **(E)** C-terminal extension lengths. Standard deviation shaded red behind the mean. **(F)** Comparison of hydrophobicity between consensus sequences of Type 1 (left) and Type 2 (right) McdB proteins. **(G)** Table quantifying biased amino acid compositions among McdB protein domains. **(H)** PONDR disorder scatter plot for all Type 1 (red) and Type 2 (blue) McdB proteins. **(I)** PONDER disorder plot between consensus sequences of Type 1 and Type 2 McdB proteins.

Consistent with the low hydrophobicity of McdB proteins, PONDR predicted that McdB proteins were largely disordered **(Figure 4H)**. Indeed, we found that the mean disorder of McdB proteins was 64%; PONDR predicted that some McdB proteins were as high as 96% disordered **(Figure 4H)**. Interestingly, while the consensus sequence for Type 1 McdB’s was predicted to be largely ordered in a central region that partially corresponded to a hydrophobic patch and that the bulk of disorder was positioned towards the N-terminal region of the proteins **(Figure 4FI)**, the consensus sequence for Type 2 McdB’s was predicted to possess much greater disorder in the N- and C-terminal extensions from the central coiled-coils and predicted to be ordered in a region that corresponded to the coiled-coils **(Figure 4I)**. Collectively, these features were largely responsible for the diversity among McdB proteins.

### McdB Undergoes Liquid-Liquid Phase Separation in vitro

The shared hallmarks of McdB proteins include: (i) large regions of low-complexity that greatly vary in length, (ii) intrinsically disordered regions, (iii) repetitive and biased amino acid compositions, (iv) low hydrophobicity, and (v) extreme multivalency. All are features that are characteristic of proteins shown to undergo Liquid-liquid phase separation (LLPS) **(Kato et al., 2012; Oldfield and Dunker., 2014; Elbaum-Garfinkle et al., 2015; Lin et al., 2015; Molliex et al., 2015; Nott et al., 2015; Varadi et al., 2015; Jain et al., 2016)**. LLPS refers to the ability of an otherwise homogeneous solution of macromolecules (ex. proteins or nucleic acids) to spontaneously demix into a dilute phase and dense phase that resembles water droplets **(Figure 5A) (Reviewed in: Alberti et al., 2019)**. The two liquid-like phases coexist and can in some cases undergo further reversible phase transitions to form gels and solids depending on solution conditions (i.e. macromolecule concentration, pH, salt type and concentration, and temperature); some transitions are irreversible under physiological conditions, such as amyloid-like fibers **(Alberti et al., 2019)**. This phenomenon has received increased attention in cell biology due to the discovery that this process accounts for the compartmentalization of biological functions through the formation of membraneless organelles, also recently termed biomolecular condensates **(Shin and Brangwynne, 2017).**

**Figure 5:**
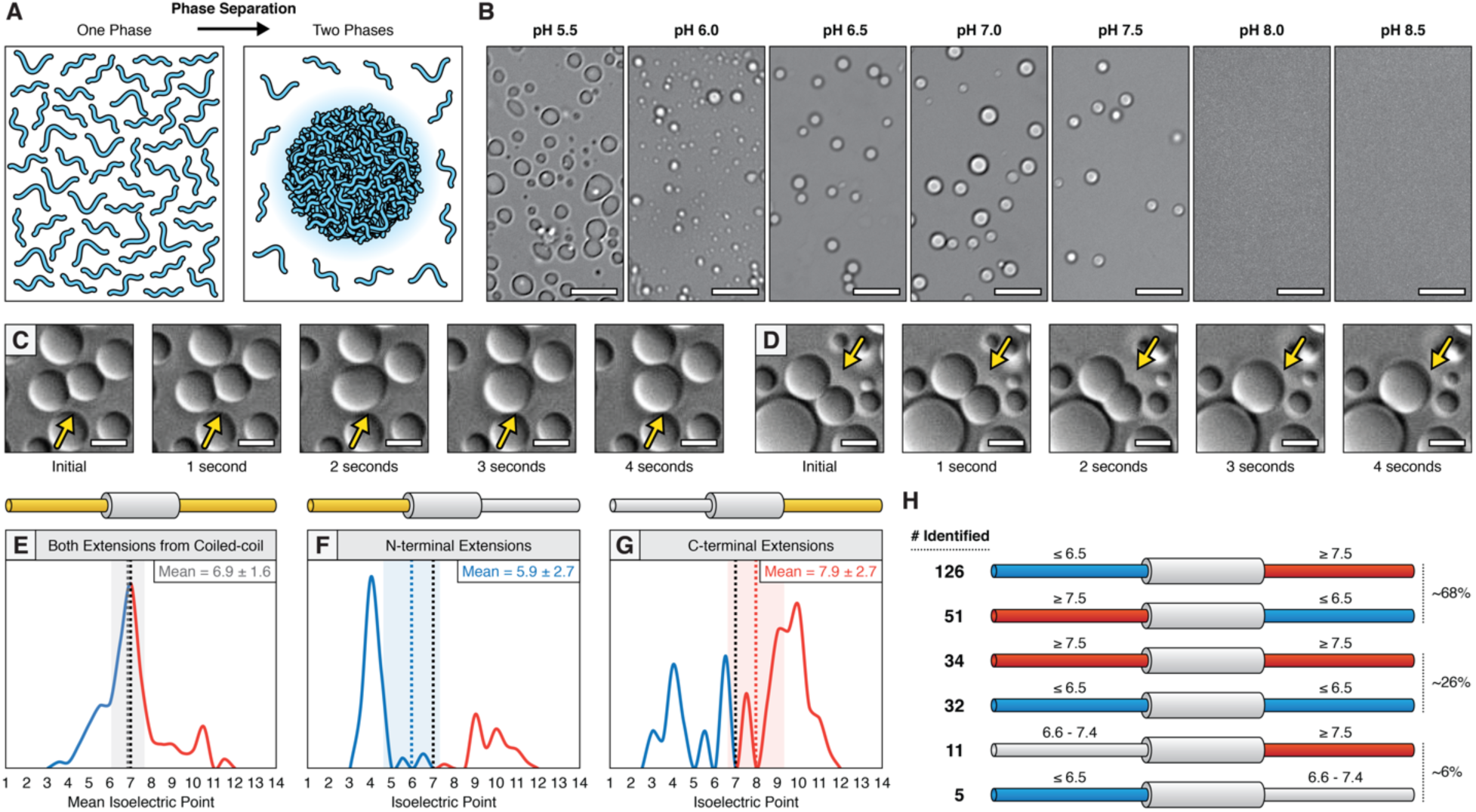
*S. elongatus* McdB undergoes liquid-liquid phase separation. **(A)** Cartoon illustration of protein liquid-liquid phase separation from one phase to two phases. **(B)** Microscopy images of *S. elongatus* McdB droplets under varying pH. Scale bar = 10 *µ*m. **(C**,**D)** McdB droplet fusion events (yellow arrows). Scale bar = 5 *µ*m. **(E)** Plot of the isoelectric point for both N- and C-terminal extensions of Type 2 McdB proteins. **(F)** Plot of the isoelectric point for the N-terminal extensions of Type 2 McdB proteins. **(G)** Plot of the isoelectric point for the C-terminal extensions of Type 2 McdB proteins. **(H)** Illustration of the number of Type 2 McdB proteins identified that have patterned charge distributions. Isoelectric point range listed above each extension.

Given the general features of the McdB proteins we identified in this study **(Figure 4)**, and that the bulk of our biochemical characterization thus far on the McdAB system has been performed with *S. elongatus*, we next wanted to test the hypothesis that *S. elongatus* McdB could undergo phase separation. We expressed and purified *S. elongatus* McdB fused at its N-terminus with a 6x-histidine tag for affinity purification and a SUMO tag to form His-SUMO-McdB. A SUMO-tag has been shown to inhibit the LLPS activity of proteins, thus simplifying the purification protocol **(Schuster et al., 2018)**. Also, cleavage of the SUMO-tag by the addition of the Ulp1 protease allows for precise, time-controlled induction of LLPS. Under physiologically relevant pH and salt concentration (pH 7.5, 150 mM KCl), native McdB displayed phase-separation upon the addition of Ulp1 **(Figure 5B)**.

The local environment of the carboxysome in *S. elongatus* cells has been suggested to be more acidic relative to the more basic cytosol because of bicarbonate accumulation prior to its conversion to CO_2_ by carbonic anhydrase **(Mangan et al., 2016)**. Given that McdB strongly colocalizes with carboxysomes *in vivo* **(Figure 3CF: MacCready et al., 2018**), we assayed the LLPS activity of McdB across a broad pH range (5.5 – 8.5) **(Figure 5B)**. We found that McdB (30 *µ*M) readily formed droplets following cleavage of the His-SUMO tag across a pH range of 5.5 – 7.5 **(Figure 5B)**. Between pH 5.5 and 6.5, droplets were less spherical in shape and did not readily fuse, which is suggestive of a gel-state **(Alberti et al., 2019)**. However, between pH 6.5 – 7.5, McdB droplets displayed liquid-like behaviors such as the fusion of two adjacent droplets into one **(Figure 5B)**. At or above pH 8, McdB did not undergo LLPS **(Figure 5B)**. Even with significantly higher concentrations of McdB (167 *µ*M), McdB still displayed liquid-like behavior with frequent fusion events **(Figure 5CD and Videos 1-2)**. Complete cleavage of the His-SUMO tag was verified across the assayed pH-range **(Figure S3)**. The data suggest that the acidic nature of the carboxysome, relative to the cytosol of *S. elongatus* at pH 8 **(Mangan et al., 2016)**, may facilitate local phase-separation of McdB in the vicinity of carboxysomes. To our knowledge, this is the first direct demonstration of LLPS behavior of a ParA partner protein.

### Charge Distribution May Contribute to McdB Phase Separation

Multivalency (charge distribution in the primary amino acid sequence) is an important feature contributing to LLPS **(Li et al., 2012; Harmon et al., 2017)**. For example, LLPS among the most well-studied systems involves interaction between positively charged residues within an intrinsically disordered region of proteins and the negatively charged backbone of DNA or RNA **(Alberti et al., 2019)**. However, we previously showed that McdB does not interact with DNA, but instead interacts with the carboxysome shell proteins CcmK2, CcmK3, CcmK4, CcmL and CcmO **(Figure 3F: MacCready et al., 2018)**. We asked where the multivalency within our system could arise. While shell proteins have surface accessible acidic and basic residue patches that McdB could interact with, shell proteins are not intrinsically disordered nor are the acidic and basic regions of all shell proteins that McdB interacts with similar **(Sommer et al., 2017)**. Therefore, given that *S. elongatus* McdB could robustly phase separate on its own, without the addition of any interacting partner (i.e. a carboxysome shell component), we next set out to determine whether Type 1 and Type 2 McdB proteins were intrinsically multivalent.

Analysis of the single N-terminal extension from the C-terminal coiled-coil of *S. elongatus* Type 1 McdB revealed that the isoelectric point of roughly the first half of the extension was ∼8.5 (amino acids 1 to 50) and the second half was ∼4.3 (amino acids 51 to 108). For Type 2 McdB proteins that have N- and C-terminal extensions from a central coiled-coil, the average isoelectric point of both extensions combined was essentially neutral (6.9 ± 1.6) **(Figure 5E)**. Intriguingly however, we found that the average isoelectric point of the vast majority of N- and C-terminal extensions were either negatively- or positively-charged **(Figure 5F-G)**. Moreover, when looking at Type 2 McdB proteins individually, we found that the vast majority (∼68%) had extremely inverted charges distributed between their N- and C-terminal extensions; ∼48% had a negatively charged N-terminus and positively charged C-terminus, and ∼20% had a positively charged N-terminus and negatively-charged C-terminus **(Figure 5H and Figure S4)**. Roughly 26% of McdB proteins had both extensions share a similar charge with 13% having both positively-charged extensions, and ∼13% having both negatively-charged extensions **(Figure 5H and Figure S4)**. Lastly, only ∼6% of McdB proteins had one charged extension and one neutral extension; ∼4% had a neutral N-terminus and positively-charged C-terminus and ∼2% had a negatively-charged N-terminus and neutral C-terminus **(Figure 5H and Figure S4)**. Together, these results suggest that McdB is a polyampholyte with biphasic charge distributions within intrinsically disordered regions that may contribute to its LLPS activity.

## Discussion

### McdAB systems are widespread among β-cyanobacteria

Carboxysomes are essential protein-based organelles in cyanobacteria. To ensure that each daughter cell receives an optimum quantity of carboxysomes following cell division, the McdAB system equidistantly positions each carboxysome relative to one another throughout the cell **(Figure 1C)**. In our previous study, we were unable to determine how widespread the McdAB system was among cyanobacteria. While we were able to identify one McdAB system outside of *S. elongatus*, in *Gloeobacter kilaueensis* JS1, and show that its McdB protein was capable of loading onto carboxysomes in *S. elongatus*, it was not obvious why BLASTp results for *S. elongatus* McdAB were so poor and why *G. kilaueensis* JS1 McdAB were so different than those from *S. elongatus* (22.5% and 18.4% pairwise identity to *S. elongatus* McdA/B) **(Figure 8B; MacCready et al., 2018)**.

Since McdAB were situated near the carboxysome operon in *G. kilaueensis* JS1, we reasoned that neighborhood analysis was a better method to identify McdAB proteins in other cyanobacteria. Indeed, we were able to identify 56 McdA and 75 McdB sequences that clustered near carboxysome components throughout cyanobacteria **(Figure 1E and Table 1)**. Given this much larger sample size of McdAB proteins, we were able to identify highly-conserved regions and features that permitted further identification of McdAB sequences that did not cluster near carboxysome components in other cyanobacteria. In total, we identified 248 McdA and 285 McdB sequences, showing that the McdAB system is widespread among β-cyanobacteria **(Figure 2A)**.

These results have broader implications for understanding carboxysome positioning in other species. For example, cyanobacteria can display a wide range of morphologies from unicellular to specialized multicellular **(Figure 2C)**. While carboxysomes are linearly-spaced or hexagonally-packed along the long axis of rod-shaped *S. elongatus* cells, our prior modeling suggested that McdAB positioning is influenced by cellular geometry, but still operates within spherical cyanobacterial cells to optimally space carboxysomes from one another **(Figure 7E; MacCready el at., 2018)**. Likewise, many heterocystous and ramified cyanobacteria, while appearing filamentous, are a chain of connected smaller cells that individually display more rounded morphologies. Transmission electron micrographs from heterocystous cyanobacteria reveal that the McdAB system likely hexagonally packs carboxysomes **(Montgomery et al., 2015)**. Therefore, given the ubiquity of McdA/B across these morphologies **(Figure 2C)**, an understanding of how this system behaves within these unique cellular geometries is of profound interest.

Lastly, we found that a large clade of cyanobacteria on the right side of our phylogenic tree appeared to lack the McdAB system **(Figure 3)**. Many of these species, including *Myxosarcina sp.* GI1, *Xenococcus sp.* PCC 7305, *Stanieria sp.* NIES-3757, *Stanieria cyanosphaera* PCC 7437, *Pleurocapsa sp.* CCALA 161, and *Pleurocapsa sp.* PCC 7319 display baeocystous morphologies and some species, including *Spirulina major* and *Spirulina subsalsa*, while filamentous, have extremely spiralized morphologies. One possible reason for the absence of the McdAB system in these cellular morphologies is that their chromosome organization, cell growth, and division strategies may be incompatible.

### Two types of McdAB systems exist in β-cyanobacteria

Our analysis has identified two distinct McdAB systems in β-cyanobacteria. Type 2 systems are by far the most represented among cyanobacteria (> 98% of species). This finding in particular explains why BLASTp results for the Type 1 system of *S. elongatus* McdAB were so poor. There are several features that distinguish Type 1 and Type 2 McdAB systems. For example, Type 2 McdA proteins possess the “signature lysine” residue in the Walker-A box ATP-binding motif; a feature that defines this family of ATPases as “ParA-like”. This result was intriguing given that Type 1 McdA proteins, like that of *S. elongatus*, lack this signature lysine in the Walker A box. Instead, Type 1 McdA proteins use a lysine residue (K151) distantly located on the C-terminal half of the protein to fulfill this role **(Schumacher et al., 2019) (Figure 2D)**. Also exclusive to Type 1 McdA proteins is a large mid-protein extension adjacent to the Walker-B box. Overall, it is the Type 2 McdA protein that bears the closest similarity to ParA-family ATPases.

The differences in amino acid sequence in and around the ATP-binding motifs suggest these two McdA types may also have differences in ATPase activity. Indeed, we previously found that the Type 1 McdA of *S. elongatus* has voracious ATPase activity and a relatively-low ATPase stimulation by McdB **(Figure 2GH: MacCready et al., 2018)**. This is in stark contrast to ParA family members involved in bacterial DNA segregation, which have feeble intrinsic ATPase activities, but are strongly stimulated by their cognate ParB protein. Therefore, it will be interesting to compare the ATPase activities of the two McdA types, as well as the stimulatory activities of their respective McdB proteins. How Type 1 and Type 2 systems differ mechanistically, and whether these differences correlate to known factors influencing carboxysome positioning such as carboxysome size and quantity **(Gonzalez-Esquer et al., 2016; MacCready et al., 2018)**, genome copy number and volume **(Griese et al., 2011)**, redox state of cells **(Sun et al., 2016, 2019)**, and McdAB stoichiometry **(MacCready et al., 2018)**, as well as unknown factors including McdB LLPS behavior and/or intracellular salt/pH conditions (see below), is important for understanding the diversity and evolution of the McdAB system among cyanobacteria.

While McdA proteins differed in certain regions, they were largely conserved compared to McdB proteins that displayed extreme diversity. The lack in conservation is presumably a result of relaxed selection, since any two closely related cyanobacterial species had McdB proteins that poorly aligned **(Figure S2B)**. Despite this lack in conservation, we were able to identify several shared features among McdB proteins. All possessed a highly charged N-terminus, a glutamine-rich region towards the center of the protein, a coiled-coil domain, and an invariant tryptophan residue within the last four amino acids of the protein **(Figure 2E)**. For several ParA family ATPases, the N-terminus of the partner protein is necessary and sufficient for stimulation of ParA ATPase activity **(Radnedge et al., 1998; Ravin et al., 2003; Barillà et al., 2007; Ah-Seng et al., 2009)**. Where tested, a critical N-terminal arginine or lysine residue stimulates the ATPase activity of the cognate ParA. The charged N-terminal domains of Type 1 versus Type 2 McdB proteins appeared inverted (location of positively- and negatively-charged residues) **(Figure 2E)**. This finding suggests a possible difference in how Type 1 and Type 2 McdB proteins interact with their McdA partner, potentially through this N-terminal region. The central glutamine-rich region likely plays a role in LLPS activity of McdB proteins (see below). A striking difference between the two McdB types was the placement of the coiled-coil. Type 1 McdB proteins have a coiled-coil positioned at their C-terminus, whereas the coiled-coil of Type 2 McdB proteins was centrally located **(Figure S4)**. Coiled-coils provide an interface for stable oligomerization into defined higher-order species **(Apostolovic et al., 2010)**. Consistent with our sequence-based predictions, a recently solved structure of the coiled-coil domain in a Type 1 McdB-like protein showed stable dimerization in an antiparallel fashion **(Schumacher et al., 2019)**. As for the invariant tryptophan at the C-terminus of all McdBs, it is intriguing that many proteins involved in the assembly of viral- or phage-capsids also encode for a tryptophan residue at their C-terminus **(Deeb, 1973; Tsuboi, et al., 2003; Komla-Soukha and Sureau, 2006; Marintcheva et al., 2006)**. Given the capsid-like icosahedral structure of the carboxysome, it is attractive to speculate that the C-terminal tryptophan of McdB is involved in its association with the carboxysome shell.

### The McdAB System is not present in α-cyanobacteria

One of our most striking findings was that the McdAB system was completely absent in α-cyanobacteria. While there are several differences between α- and β-carboxysomes, we expected this difference would largely be reflected in the putative carboxysome interaction domains of McdB proteins. We did not anticipate a complete absence of the system. Even more surprisingly, we found that not just McdA, but the entire ParA family of ATPases were significantly underrepresented among α-cyanobacteria. α-cyanobacteria lack plasmids and have small genomes (most *Prochlorococcus* genomes are smaller than 2 Mb) **(Scanlan et al., 2009)**. Therefore, one explanation for the lack of ParA-type ATPases is that many canonical partitioning systems cannot function in α-cyanobacteria. Also, the absence of plasmids in *α*-cyanobacteria limits one of the sources of horizontal gene transfer for inheriting the McdAB system.

Alternative positioning mechanisms may exist for α-carboxysomes in α-cyanobacteria. For example, α-carboxysomes are known to tightly interact with polyphosphate bodies **(Iancu et al., 2009)**. If polyphosphate bodies are strategically positioned in α-cyanobacteria, and α-carboxysomes tightly interact with these structures, this interaction would provide a “pilot-fish” mechanism by which both are equidistantly positioned throughout cells prior to cell division; making the McdAB system unnecessary. Another possibility is that α-carboxysomes are actively positioned by a polymer-based system, such as the actin-like ATPase MamK that mediates magnetosome positioning **(Toro-Nahuelpan et al., 2016)** or the tubulin-like GTPase TubZ that positions viral DNA **(Oliva et al., 2012)**.

Since α-carboxysomes are proposed to have originated in Proteobacteria and were horizontally transferred into cyanobacteria **(Marin et al., 2007; Rae et al., 2013)**, it was surprising that the McdAB system was absent in α-cyanobacteria largely because many carboxysome operons of Proteobacteria encode a ParA-type ATPase followed by a small coding sequence. Whether this ParA-type ATPase and the small downstream coding sequence constitutes the McdAB system for *α*-carboxysomes in Proteobacteria is of great interest for understanding the evolution, diversity, and mechanisms of carboxysome positioning across the bacterial world.

### The S. elongatus McdAB system might have a unique origin

The phylogenic placement of *S. elongatus* and Type 1 McdAB systems relative to Type 2 systems suggest the latter is more ancestral. *G. kilaueensis* JS1 is one of the most primitive species among living cyanobacteria due to the lack of thylakoid membranes **(Rippka et al., 1974; Guglielmi et al., 1981; Nakamura et al., 2003)**. It possesses the widely-distributed Type 2 system, suggesting that the Type 1 system of *S. elongatus* was acquired more recently. We obtained strong bootstrap support for *S. elongatus* sharing an immediate common ancestor with α-cyanobacteria, which lack the McdAB system, and four cyanobacteria where the McdA/B system could not be identified **(Figure 3)**. Many Type 1 McdA homologs identified here by BLASTp are plasmid encoded (the sole ParA-type ATPase on the plasmid) and the N-terminal domain of Type 1 McdB’s are nearly identical to the small coding sequences following these plasmid-encoded McdA-like proteins. Therefore, the most parsimonious hypothesis is that the plasmid encoded ParA-type system used to partition plasmids was horizontally transferred into *S. elongatus*, genomically integrated, and evolved to position carboxysomes. A similar evolutionary path would explain the positioning of a diversity of cargos by ParA family ATPases **(Lutkenhaus, 2012; Keikebusch and Thanbichler, 2014; Vecchiarelli et al., 2012)**.

Since the Type 2 system was present at the earliest known point of cyanobacterial evolution, it is not clear why the Type 1 system would have had a selective advantage over the Type 2 system, presumably present in the ancestors of *S. elongatus*. It is also interesting that we were unable to identify the McdAB system in several modern cousins of *S. elongatus* **(Figure 3)**. An alternative hypothesis could be that this clade of *β*-cyanobacteria (which now includes modern *S. elongatus*) lost the Type 2 system, similar to the *β*-cyanobacterial clades found on the right side of our tree, and that *S. elongatus* then acquired the Type 1 system. As more cyanobacterial genomes are sequenced, how the Type 1 McdAB system evolved will become more apparent.

### McdB is a phase separating protein

Our finding that *S. elongatus* McdB undergoes phase separation *in vitro* is intriguing on multiple fronts. First, two recent studies have demonstrated that α- and β-carboxysome formation involves LLPS **(Oltrogge et al., 2019; Wang et al., 2019)**. With β-carboxysomes, the protein CcmM aggregates RuBisCO to form a procarboxysome **(Cameron et al., 2013)**. CcmM exists in two forms: (i) full-length CcmM (58 kDa), which contains a carbonic anhydrase-like domain followed by three RuBisCO small subunit-like domains (SSLD) separated by flexible linkers, and (ii) short-form CcmM (35 kDa), which lacks the carbonic anhydrase-like domain **(Long et al., 2010)**. Binding of SSLD regions of CcmM between RbcL dimers scaffolds RuBisCO molecules and induces LLPS **(Wang et al., 2019)**. Likewise, in α-carboxysomes of the chemoautotrophic proteobacterium *Halothiobacillus neapolitanus*, N-terminal repeated motifs within the intrinsically disordered protein CsoS2 mediates interaction with RuBisCO and induces LLPS **(Oltrogge et al., 2019)**. The pyrenoid does not have a protein shell, but serves as the functional equivalent to carboxysomes in eukaryotic algae. One study of the pyrenoid showed that its RuBisCO matrix also exhibits liquid-like properties when interacting with the intrinsically disordered protein EPYC1 in *Chlamydomonas reinhardtii* **(Freeman Rosenzweig et al., 2017)**. Collectively, these studies suggest that LLPS is a common feature underlying the formation of a RuBisCO matrix through interactions with intrinsically disordered proteins. Given that McdB is also an intrinsically disordered protein that undergoes LLPS **(Figure 5BCD and Videos 1-2)** and our previous data that increased levels of McdB drastically increase carboxysome size **(Figure 5M and Figure S5AG: MacCready et al., 2018)**, it is intriguing to speculate that the LLPS activities of McdB and the carboxysome core are related and potentially influence each other. It is possible that the McdAB system not only positions carboxysomes in the cell, but also maintains homeostasis of a liquid-like carboxysome core.

Our results that *S. elongatus* McdB formed droplets across a broad pH range is informative. It has been previously suggested that a pH gradient exists between the cytosol and carboxysomes **(Menon et al., 2010; Whitehead et al., 2014)**. Experimental evidence in *S. elongatus* showed that while the pH of the cytosol of *S. elongatus* is ∼8.4 under light conditions, the carboxysome is predicted to be more acidic (pH ∼6.0-7.0) **(Mangan et al., 2016)**. An acidic carboxysome has been suggested to increase the maximum carboxylation rate of RuBisCO and reduce the amount of HCO_3_-uptake required to saturate RuBisCO **(Mangan et al., 2016)**. Our *in vitro* data suggests that McdB would be soluble in the cytosol (pH ≥ 8) **(Figure 5B)** and undergo LLPS at or near acidic carboxysomes (pH ≤ 7.5) **(Figure 5B)**. However, dark adapted cells of *S. elongatus* have a cytosolic pH ∼7.3 **(Mangan et al., 2016)**, so McdB behavior might differ in cells under light and dark conditions. Moreover, cyanobacteria possess a biological circadian clock that precisely operates on the 24-hour rotational period of the earth. Circadian rhythms primarily enable cyanobacteria to anticipate, adapt and respond to daily light cycles by translating environmental cues into changes in gene expression **(Reviewed in: Cohen and Golden, 2015)**. In *S. elongatus*, oscillatory patterns of gene expression are driven by phosphorylation of the master output transcriptional regulator protein RpaA. Phosphorylation of RpaA has previously been shown to bind ∼170 promoters of the *S. elongatus* chromosome **(Markson et al., 2013)**; one of which is the promoter for the *mcdAB* operon. Therefore, it will be interesting to explore the role of McdB LLPS activity at carboxysomes, and how circadian rhythms and light-dark conditions influence McdAB expression, dynamics, and function.

Lastly, McdB is the first example of a ParA partner protein shown to have LLPS activity. It is attractive to speculate that under appropriate conditions, other ParA partner proteins exhibit similar LLPS behaviors that are critical to their *in vivo* function. The most obvious examples are ParB proteins that bind to a centromere-like DNA sequence called *parS* and are involved in plasmid and chromosome segregation in bacteria **(Reviewed in Wang et al., 2013; Baxter and Funnell, 2014; Bouet et al., 2014)**. Through ChIP approaches, it is well known that thousands of ParB dimers associate with broad regions of DNA adjacent to the *parS* site, a phenomenon known as “spreading” **(Rodionov et al., 1999; Murray et al., 2006; Breier and Grossman, 2007; Sanchez et al., 2015; Debaugny et al., 2018)**. However, the actual structure of the ParB-DNA mega-complex remains unclear. *In vivo*, fluorescent fusions of ParB form a massive punctate focus at the location of a *parS* site on the chromosome or plasmid. These ParB foci segregate after DNA replication in a ParA-dependent manner and have also been shown to undergo rapid fission/fusion events, whereby ParB foci segregate and then snap back together **(Sengupta et la., 2010)**. FRAP measurements of ParB exchange at these foci also suggest these complexes are highly dynamic **(Debaugny et al., 2018)**. Together, the data are consistent with ParB forming a biomolecular condensate with its *parS* site and its adjacent DNA. LLPS activity of ParB proteins on *parS*-containing DNA substrates has yet to be directly observed *in vitro*. However, *in silico* models have proposed that the currently known affinities of ParB-ParB and ParB-DNA associations could induce the formation of a condensate **(Broedersz et al., 2014; Sanchez et al., 2015; Debaugny et al., 2018)**. The vast majority of proteins capable of LLPS do so by interacting with DNA/RNA **(Reviewed in: Alberti et al., 2019)**. McdB however does not bind DNA and is capable of LLPS activity on its own. A future direction of research is aimed at understanding how the LLPS activity of McdB, and other ParA partner proteins, is involved in the spatial organization of their cognate cargos.

### McdB charge distribution might contribute to LLPS

McdB proteins across evolutionary time possess many of the features that enable LLPS including intrinsic disorder, low hydrophobicity, biased amino acid compositions, and extreme multivalency **(Reviewed in: Alberti et al., 2019) (Figure 4)**. When analyzing the multivalent properties of McdB proteins, we found that the vast majority were polyampholytes with biphasic charge distributions between their N- and C-terminal extensions flanking the coiled-coil domain **(Figure 5H and Figure S4)**. The reason for such a shared feature is not obvious. While genetic drift possibly accounts for the intrinsic disorder and lack of primary sequence conservation among McdB proteins, it is unlikely that genetic drift alone could account for the patterned charge distributions. Indeed, given that the vast majority of McdB proteins exhibit an inverted charged patterning along their N- and C-terminal extensions **(∼68%, Figure 5H and S4)**, a more parsimonious explanation could be that the charged extensions are important for McdB associations with itself or with carboxysome shell proteins. While not a true McdB protein, the structure of the coiled-coil domain of a plasmid-encoded McdB-like protein from the cyanobacterium *Cyanothece sp.* PCC 7424 (PCC7424_5530) demonstrated an antiparallel association to form a dimer **(Schumacher et al., 2019)**. Parallel or antiparallel assembly of the coiled-coil domains of McdB would either align like- or oppositely-charged extensions. The role of multivalency, charge patterning, and dimerization on the LLPS activity of McdB are interesting avenues of future investigation.

### mcdA and mcdB genes often cluster with the minor carboxysome shell genes ccmK3 and ccmK4

Of the *mcdA* and *mcdB* genes we found near carboxysome components, the majority clustered near the minor shell genes *ccmK3* and *ccmK4* **(Figure 1E and 2A)**; two shell genes often found together, but at a different locus than the carboxysome operon **(Sommer at al., 2017)**. The finding is consistent with our prior results showing that McdB interacts with CcmK3 and CcmK4 in a bacterial two-hybrid system **(Figure 3F: MacCready et al., 2018)**. Moreover, carboxysomes have been shown to cluster following the deletion of CcmK3 and CcmK4 in a manner that is reminiscent of carboxysome aggregation in our *ΔmcdB* strain **(Rae et al., 2013; Figure 5J: MacCready et al., 2018)**. How McdB and CcmK3/CcmK4 interact is still an open question. A recent study showed that CcmK3 and CcmK4 can form homohexamers, as well as heterohexamers that further assemble into dodecamers under certain conditions **(Sommer et al., 2019)**. Metabolite channeling into and out of carboxysomes is believed to occur via the central pores of these hexameric shell proteins **(Dou et al., 2008; Kinney et al., 2011)**. Since the pore formed by CcmK4 homohexamers differs from those formed by CcmK3/CcmK4 heterohexamers, it is proposed that homo- and hetero-hexamic species alter carboxysome permeability, thereby modulating metabolite channeling across the protein shell **(Sommer et al., 2019)**. Interestingly, CcmK3/CcmK4 dodecamer formation was found to be influenced by pH (relative dodecamer abundance of 36% at pH 7.0 and 3% at pH 8) **(Sommer et al., 2019)**. Likewise, we find here that McdB LLPS behavior is also greatly influenced by transitions in this pH range; soluble at a pH ≥ 8 and formed liquid droplets at a pH ≤ 7.5 **(Figure 5B)**. Placing these findings in the context of current models that propose the carboxysome as more acidic (pH ∼ 7.0) than the cytosol (pH ∼ 8.4) in *S. elongatus* **(Menon et al., 2010; Whitehead et al., 2014; Mangan et al., 2016)**, several important questions are revealed. How does McdB interact with CcmK3/CcmK4 assemblies? Is this interaction influenced by McdB LLPS activity and by the differential pH of the carboxysome versus the cytoplasm? Finally, does McdB influence metabolite channeling into and out of carboxysomes and is this activity dependent on LLPS behavior of McdB? It is attractive to speculate that local pH changes due to the metabolic activities of the carboxysomes also serves to regulate its composition, structure, and function. Together, our results have broad implication for understanding the diversity of the McdAB system, the mechanisms of carboxysome positioning among carbon-fixing bacteria, and the role of LLPS in the biogenesis and spatial organization of carboxysomes.

## Acknowledgements

We would like to thank Lindsay Matthews from the Simmons lab for providing the Ulp1 enzyme. This work was supported by the National Science Foundation (Award Number 095064), research initiation funds provided by the MCDB Department to AGV, University of Michigan, and by research funds from the Michigan Life Sciences Fellowship program to JSM.

## Author Contributions

JSM. and AGV. conceived the project. JSM., JLB., and AGV., designed experiments. JSM. and JLB. performed all experiments. JSM and AGV wrote the manuscript. All authors discussed results and edited the manuscript.

## Supplemental Information

**Figure S1:**
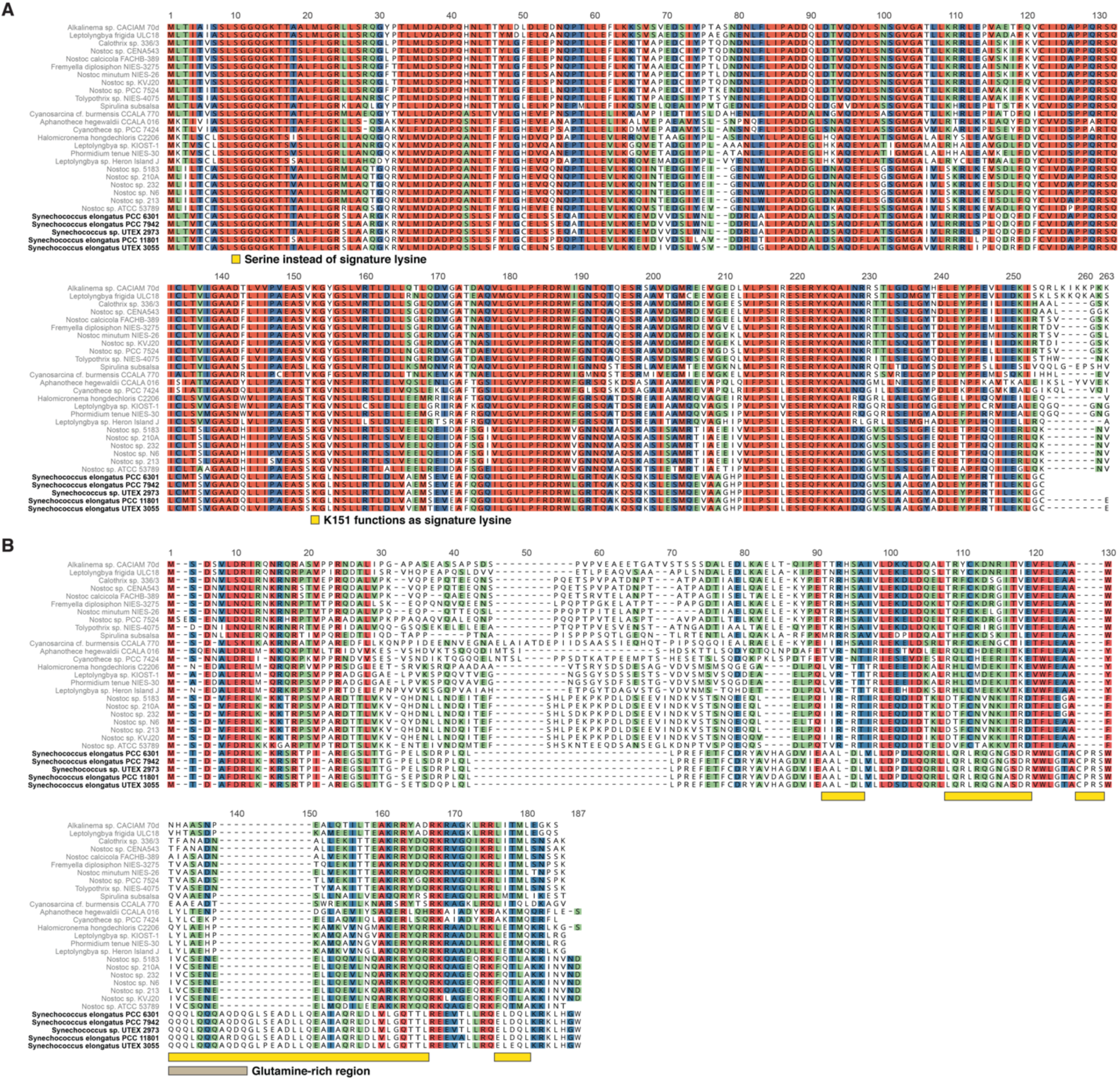
BLASTp results for S. elongatus McdA and McdB. **(A)** Multiple sequence alignment of BlastP hits for *S. elongatus* McdA. Serine and lysine residues distinguishing these proteins from classic ParA proteins (yellow box below alignment) **(B)** Multiple sequence alignment of BlastP hits for *S. elongatus* McdB. Regions of dissimilarity between *S. elongatus* McdB and the protein products of the downstream open reading frame that follow the McdA-like proteins from panel **A** (yellow boxes below alignment). Glutamine-rich region from S. elongatus McdB (brown box below alignment).

**Figure S2:**
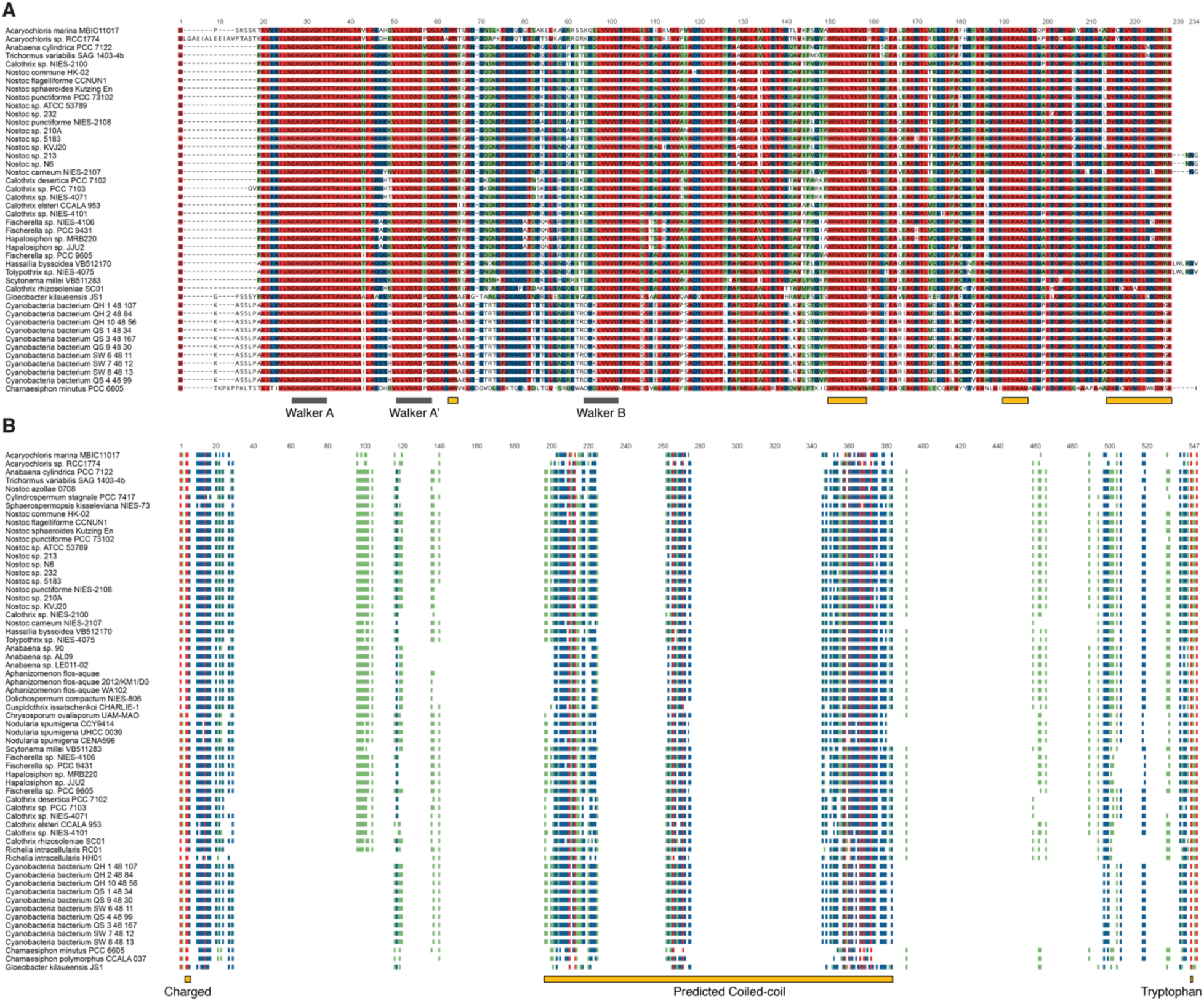
Conserved features among McdA and McdB proteins near carboxysome-related components. **(A)** Multiple sequence alignment of McdA proteins found near carboxysome-related components. Domains conserved with classic ParA proteins (grey boxes below alignment). Domains unique to identified McdA proteins (yellow boxes). **(B)** Multiple sequence alignment of McdB-like proteins found near carboxysome-related components. The conserved features of McdB-like proteins: Charged N-terminal domain, predicted central-coiled coil and C-terminal tryptophan are highlighted (yellow boxes).

**Figure S3:**
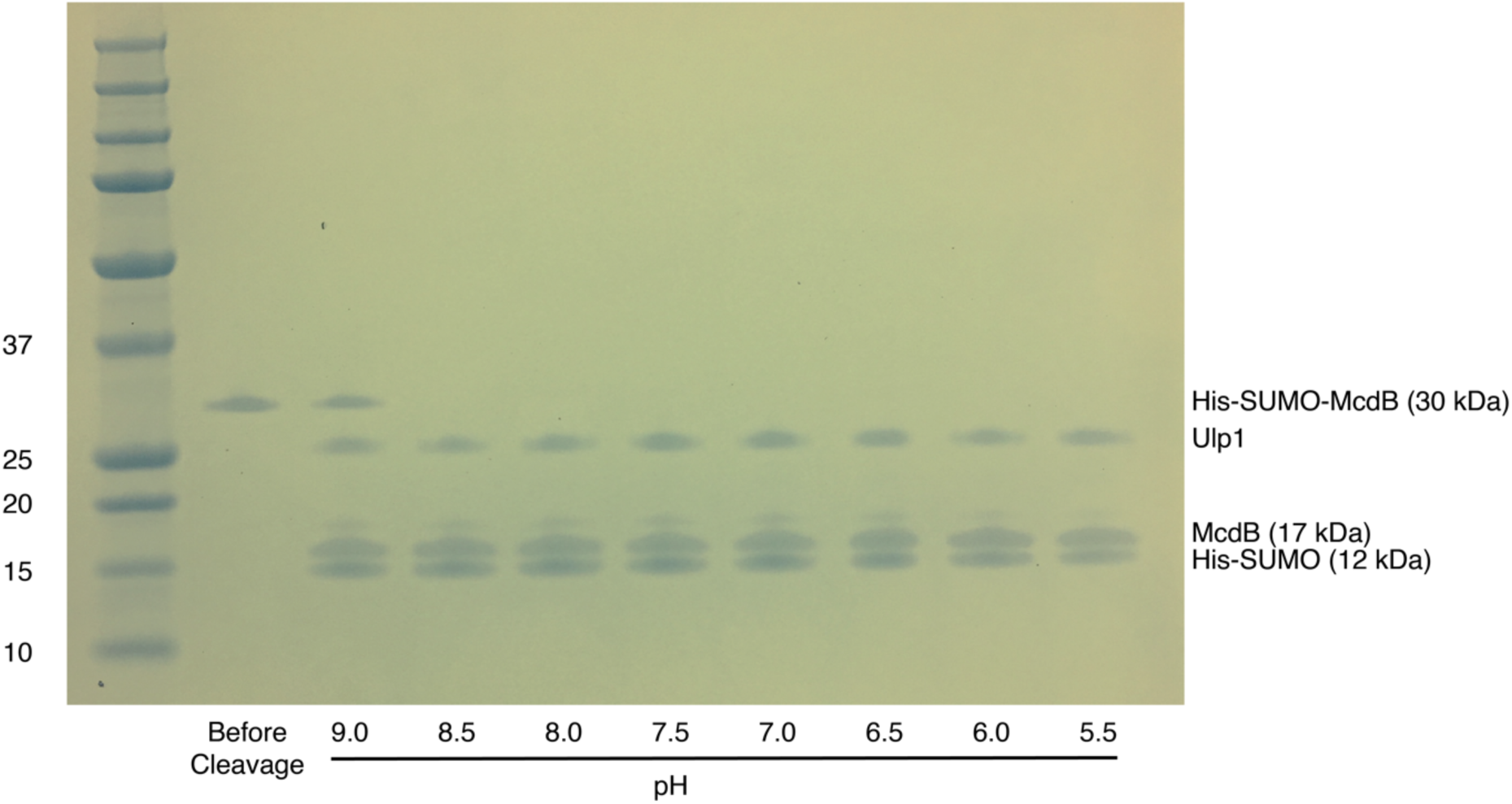
*S. elongatus* His-SUMO-McdB is completely cleaved by Ulp1 across the pH range ranged tested for McdB LLPS activity. Coomassie-stained SDS-PAGE shows that His-SUMO-McdB is completely cleaved following Ulp1 digestion within the pH range of 5.5 – 8.5. Complete cleavage was not achieved at pH 9.

**Figure S4:**
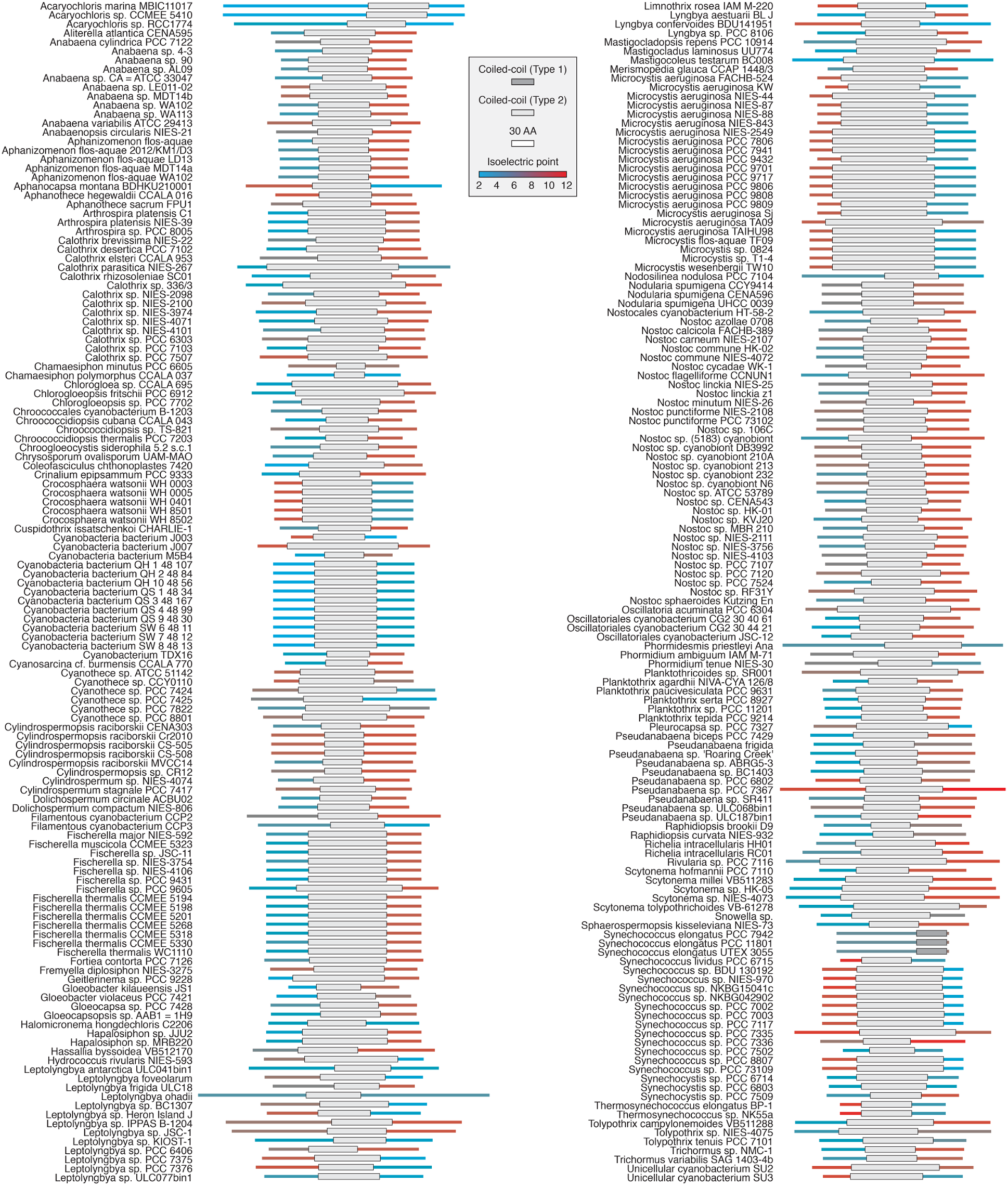
Biphasic charge distributions are a common feature of Type 2 McdB proteins. Illustration of charge distributions in the extensions among all McdB proteins. Scale bar = 30 amino acids. Coiled coils of Type 1 and Type 2 McdB proteins are dark and light grey, respectively.

## Methods

### McdAB homolog search and neighborhood analysis

Initial searches for *S. elongatus* McdA (Synpcc7942_1833) and McdB (Synpcc7942_1834) homologs were performed via BLASTp. Since BLASTp returned few hits for McdA and not a single reliable hit for McdB, we reasoned that neighborhood analysis was a better method for identifying possible homologs of these proteins. Neighborhood analysis was performed by obtaining genome assembly genbank files from NCBI for all cyanobacteria used in this study. Since genome annotations are inconsistent among cyanobacterial species and carboxysome components can be found across multiple loci, homologs of *S. elongatus* carboxysome components CcmK2 (Synpcc7942_1421), CcmK3 (Synpcc7942_0284), CcmK4 (Synpcc7942_0285), CcmL (Synpcc7942_1422), CcmM (Synpcc7942_1423), CcmN (Synpcc7942_1424), CcmO (Synpcc7942_1425), CcmP (Synpcc7942_0520), RbcS (Synpcc7942_1427), RbcL (Synpcc7942_1426), RbcX (Synpcc7942_1535), CcaA (Synpcc7942_1447) and RbcR (Synpcc7942_1980) were used as BLASTp queries to identify carboxysome components in all other cyanobacteria. We note, additional shell proteins exist among other cyanobacteria, including CcmK1, CcmK5 and CcmK6 **(Sommer et al., 2017)**. However, these proteins share high similarity to CcmK2, so they were also captured during our BLASTp search using CcmK2.

Neighborhood analysis for each carboxysome component across all cyanobacterial genomes was performed manually. We defaulted to this approach since many cyanobacterial genomes were still drafts and we wanted to determine if the contigs or scaffolds were too small to make an appropriate neighborhood determination. Moreover, we also wanted the ability to quantify the number of base-pairs and coding sequences between *mcdA* or *mcdB* and the gene of the neighboring carboxysome component. Neighborhood analysis was carried out via Biomatters Geneious v 11.1.5 by searching 10 Kbp upstream and downstream of each carboxysome component gene across all cyanobacterial species used in this study to identify coding sequences that matched our criteria for McdA or McdB. Accession numbers of identified McdA, McdB and carboxysome component(s) were recorded using Excel.

### McdA/B sequence analysis

Multiple sequence alignments for McdA proteins were performed using MAFFT 1.3.7 under the G-INS-I algorithm, whereas the E-INS-I algorithm was used for McdB proteins due to long gaps caused by intrinsic disorder. Coiled-coil predictions for McdB proteins were carried out using DeepCoil **(Ludwiczak et al., 2019)**. Predictions of disorder within McdB proteins was performed using PONDR with the VL-XT algorithm **(Romero et al. 1997; Li et al. 1999; Romero et al. 2001)**. Analysis of McdB hydrophobicity was conducted with ProtScale using the Kyte and Doolittle scale **(Kyte and Doolittle, 1982; Gasteiger et al., 2005)**.

### Phylogenetic inference

Ortholog sequences for *S. elongatus* DnaG, RplA, RplB, RplC, RplD and RplE were obtained via BLASTp for each cyanobacterium. Alignments for protein sequence was performed using MAFFT 1.3.7 under the G-INS-I algorithm and BLOSUM62 scoring matrix. The 6 resulting alignments were then concatenated into one alignment using Geneious 11.1.5. Regions of low conservation within the resulting alignment were removed using gBlocks 0.91b **(Castresana et al., 2000; Talavera et al., 2007)**. A phylogenetic tree was then estimated with maximum likelihood analyses using RAxML 8.2.11 under the LG+Gamma scoring model of amino acid substitution. Bootstrap values were calculated from 500 replicates. The resulting phylogenetic tree was then edited and visualized using iTOL3 **(Letunic and Bork, 2016)**.

### Construct development

The His-SUMO-McdB expression plasmid was generated via Gibson Assembly **(Gibson et al., 2009)** using synthetized dsDNA for His-SUMO-McdB that was inserted into a pET11b vector. Construct was verified by sequencing. Cloning was performed in *E. coli* DH5α chemically competent cells (Invitrogen).

### His-SUMO-McdB Overexpression

The His-SUMO-McdB fusion was expressed from a pET11b vector containing an ampicillin resistance cassette. All *E. coli* cultures containing the vector were grown in LB + carbenicillin (100 *µ*g/mL) at 37°C with 220 rpm shaking unless otherwise stated. The vector was transformed into competent *E. coli* BL21-AI, and a single transformant was picked to inoculate a 25 mL overnight culture. For expression, 2 L of culture was inoculated through a 1:100 dilution of the overnight culture and grown at 37°C until an OD_600_ of 0.5-0.7 was reached. Expression was induced with final concentrations of IPTG at 1mM and L-arabinose at 0.2%. Immediately after induction, cultures were cooled in an ice bath for 5 min and shifted to 18°C to grow for 16 hrs at 220 rpm. Cells were pelleted at 4,000 × g for 20 min at 4°C, flash frozen in liq. N_2_, and stored at −80°C.

### His-SUMO-McdB Purification

The following buffers were used to purify the His-SUMO-McdB fusion: lysis buffer (50 mM HEPES pH 7.6; 1 M KCl; 5 mM MgCl2; 10% glycerol; 2 mM BME; 20 mM imidazole; 0.05 mg/mL lysozyme; 0.05 *µ*L/mL benzonase; protease inhibitor), Ni buffer A (50 mM HEPES pH 7.6; 1 M KCl; 5 mM MgCl2; 10% glycerol; 2 mM BME; 20 mM imidazole), and Ni buffer B (50 mM HEPES pH 7.6; 1 M KCl; 5 mM MgCl2; 10% glycerol; 2 mM BME; 500 mM imidazole). The 2 L cell pellet was resuspended in 150 mL lysis buffer and lysed using a microfluidizer at 18,000 psi equilibrated in Ni buffer A. The lysate was centrifuged at 30,000 × g at 4°C for 30 min. The clarified lysate was decanted, syringe filtered (0.2 μm), and loaded onto a 5 mL HP HisTRAP column equilibrated in Ni buffer A. The column was washed with 25 mL Ni buffer A and then 25 mL 5% Ni buffer B. Elution was done using a 5-100% gradient of Ni buffer B while collecting 2 mL fractions for a total of 60 mL. Peak fractions were flash frozen in N_2_ (liq.) and stored at −80°C.

### Microscopy of in vitro McdB phase separation

Prior to imaging, His-SUMO-McdB samples were buffer exchanged into Ulp1 reaction buffer (150 mM KCl; 25 mM HEPES; 2 mM BME) adjusted to the indicated pH. Buffer exchange was performed using 7 K MWCO, 5 mL Zeba Spin Desalting Columns (Thermo-Fischer). All imaging was performed using 16 well CultureWells (Grace BioLabs). Wells were passivated by overnight incubation in 5% (w/v) Pluronic acid (Thermo-Fischer), and washed thoroughly with Ulp1 buffer prior to use. For cleavage experiments, 1 *µ*L of purified Ulp1 was added to 50 *µ*L of His-SUMO-McdB at the indicated concentration, and incubated at 23 °C for 2 hrs to ensure complete cleavage. Imaging of McdB droplet formation was performed using a Nikon Ti2-E motorized inverted microscope (60x DIC objective and DIC analyzer cube) with a Transmitted LED Lamp house and a Photometrics Prime 95B Back-illuminated sCMOS Camera. Image analysis was performed using Fiji v 1.0.

